# Human pulvinar stimulation engages select cortical pathways in epilepsy

**DOI:** 10.1101/2025.03.11.642694

**Authors:** Jordan A. Bilderbeek, Nicholas M. Gregg, Maria Guadalupe Yanez-Ramos, Harvey Huang, Morgan Montoya, Peter Brunner, Jon T. Willie, Jamie J. Van Gompel, Gregory A. Worrell, Kai J. Miller, Dora Hermes

**Affiliations:** Department of Physiology and Biomedical Engineering, Mayo Clinic, Rochester MN; Department of Neurology, Mayo Clinic, Rochester, MN; Medical Scientist Training Program, Mayo Clinic, Rochester, MN; Department of Neurosurgery, Washington University School of Medicine, St. Louis, MO; National Center for Adaptive Neurotechnologies, St. Louis, MO; Department of Neurosurgery, Mayo Clinic, Rochester, MN

**Keywords:** Pulvinar, stimulation, neuromodulation, posterior quadrant epilepsy, connectivity, stereoelectroencephalography, deep brain stimulation

## Abstract

The pulvinar has been proposed as an effective neuromodulation target for patients with posterior quadrant and temporal epilepsies. However, the pulvinar has a large tissue volume, multiple subnuclei, and widespread cortical connections. It remains unknown whether electrical stimulation of distinct pulvinar subregions affects the temporal, occipital, and parietal areas differently. To address this gap, we delivered single-pulse electrical stimulation to the pulvinar and measured the resulting brain stimulation evoked potentials in twelve patients undergoing stereotactic EEG for drug-resistant epilepsy. Brain stimulation evoked potentials were parameterized across the occipital, temporal and parietal cortex. Stimulation of the lateral pulvinar elicited significant brain stimulation evoked potentials in striate and extrastriate areas that diminish as stimulation shifts towards the medial pulvinar. Conversely, stimulation of the ventral aspect of the medial pulvinar produced significant lateral temporal evoked potentials, which diminish with lateral pulvinar stimulation. We also found that stimulation of the dorsomedial pulvinar evoked significant parietal responses with limited striate/extrastriate and lateral temporal responses. These results demonstrate that electrical stimulation of specific pulvinar subregions influences distinct occipital, parietal and lateral temporal areas. Selective targeting of pulvinar subregions to maximize seizure network engagement may be essential for individualized treatment of posterior quadrant and temporal epilepsies.

## Introduction

Deep brain stimulation (DBS) is a widely used treatment for drug-resistant epilepsy. By stimulating specific thalamic nuclei, DBS aims to disrupt the generation and spread of seizures.^1^ Different thalamic nuclei are targeted for different types of epilepsy. For example, the anterior nucleus of the thalamus (ANT) is FDA-approved for focal epilepsy and is commonly used for temporal and frontal lobe epilepsies due to its placement along the Papez circuit. The centromedian nucleus, which has more diffuse connections, has shown evidence for efficacy in generalized epilepsies.^2-4^ Recently, the pulvinar has emerged as a promising therapeutic target for lateral temporal and posterior quadrant epilepsies.

The pulvinar is the largest thalamic nucleus with extensive cortical connectivity and contains several subregions. The effectiveness of pulvinar DBS for epilepsy remains unproven, but results from recent investigative studies with small patient numbers show promise.^5-10^ Given the large tissue volume, multiple subnuclei, and varied structural thalamocortical connectivity, it remains difficult to know how clinicians should target these subregions for individualized seizure network targeted neuromodulation. Moreover, even in smaller thalamic nuclei, electrode positioning is highly impactful. Studies on ANT stimulation have established that understanding both stimulation location and the connected network is crucial for achieving satisfactory clinical outcomes while limiting off-target effects.^11^ Therefore, it is vital to understand whether different lateral temporal and posterior quadrant brain areas are preferentially influenced by different pulvinar subregions. In the future, such understanding may help determine optimal electrode placement for selectively modulating lateral temporal and posterior quadrant areas.

Much of our current understanding of pulvinar connectivity hinges on non-human animal studies. Primate tracing studies have shown that lateral (PuL) and inferior pulvinar (PuI) project to striate and extrastriate areas in the occipital lobe ^12-13^. The medial pulvinar (PuM) projects to parietal, temporal, frontal, orbital, and cingulate cortices.^13-14^ The connectivity follows a gradient that begins with low-level visual connectivity ventrolateral to higher cortices dorsomedially.^12,14^ Human studies using diffusion imaging and fMRI have mirrored these connectivity profiles.

These imaging studies further delineated a dorsal-ventral distinction within the medial pulvinar, showing that dorsal PuM (dPuM) has connections with the parietal cortex, while the ventral PuM (vPuM) is connected with the temporal neocortex.^15^ Previous stimulation studies measuring effective connectivity have stimulated the medial pulvinar and report widespread influence across the temporal neocortex, parietal, frontal, and insulo-opercular cortex.^16^

Using single-pulse electrical stimulation (SPES) in 12 patients with drug-resistant epilepsy undergoing stereotactic EEG (sEEG) including pulvinar electrodes, we tested whether electrical stimulation of electrodes placed in these different pulvinar subregions selectively influences connected anatomical structures. We measured brain stimulation evoked potentials (BSEPs) across the cortex as a measure of effective connectivity. We tested whether the effective connectivity in patients with epilepsy matched anatomical reports of differential pulvinar subregions connecting to striate and extrastriate areas, lateral temporal areas, and parietal cortex.

## Materials and Methods

### Subjects and electrode localization

SPES was performed in twelve patients (**Supplemental Table 1**) implanted with sEEG during clinically indicated invasive epilepsy monitoring occurring at Mayo Clinic Rochester or Barnes Jewish Hospital St. Louis. The targeting and placement of sEEG leads were solely determined by the surgical epilepsy conference consensus recommendations and clinical team, and this study did not influence the decision to place thalamic sEEG leads. Patients had differing seizure onset zones (**Supplemental Table 1**), which reduces the potential bias of a seizure network driving responses. SPES was delivered under a Mayo Clinic and Washington University in St. Louis Institutional Review Board approved study, and all patients provided informed consent.

Electrode positions were localized based on a postoperative CT co-registered with a preoperative MRI and labeled based on anatomical atlases (**Supplementary Material, ‘Methods’ section**). We included all subjects where an sEEG contact was localized to be in the pulvinar and other sEEG contacts measured from hypothesized areas of differential connectivity: the lateral temporal cortex, including the (middle and superior temporal gyri, middle and superior temporal sulci), striate and extrastriate areas (V1, V2, V3a/b, hV4, TO1-2, LO1-2, IPS0), and parietal areas (IPS1-5).

### Single-pulse electrical stimulation

Biphasic SPES was delivered through pulvinar electrodes. Stimulation conditions varied by patient (**Supplemental Table 2)**. 12-60 pulses were delivered for each stimulation site (**Supplemental Figure 1b)**. Pulses were jittered to occur randomly every 1-5 seconds to avoid neuronal steady states.

### Thalamocortical evoked potential preprocessing and analysis

Channels and segments that showed excessive electrical noise, movement artifacts, or interictal noise were excluded from analyses (reviewed by M.M, J.A.B). Raw data was converted to a trial structure centered around the onset of electrical stimulation and corrected for DC offset. Data was re-referenced using a previously established adjusted common average referencing technique.^17^ Data was baseline corrected by subtracting the mean amplitude before stimulation.

To measure the strength of the pulvinar-cortical effective connectivity, we implemented the Canonical Response Parameterization (CRP) algorithm.^18^ CRP provides statistics for BSEPs independent of their waveform and thus allows for directly comparing different pulvinar output types (**Supplementary Material, ‘Methods’ section)**. Briefly, CRP creates a cross-projection matrix across trials for each measurement electrode using a semi-normalized dot product (**Supplemental Figure 2**). The sorted distribution of cross-projection BSEP magnitudes is then tested for statistical significance and used to determine response duration. A linear kernel PCA then calculates the canonical response (*C(t)*) from the individual trials truncated to the significant response duration. We assume that strong projections elicit highly reliable BSEPs across trials. To quantify the consistency of the BSEPs, we project the canonical response *C(t)* into each trial and calculate the coefficient of determination (R^2^), derived as the mean subtracted explained variance. The R^2^ for each measurement electrode provides an estimate of the strength of the pulvinar-cortical effective connectivity: the strength with which the pulvinar effectively influences different cortical areas. We avoid using statistics based on BSEP amplitude or root-mean-squared error as these can be impacted by electrode orientation or different brain states.^19-20^ We considered those BSEPs to be significant that exhibited a median R^2^ > 0.1, mean signal-to-noise ratio > 0.5, and a p-value < 0.05 (**Supplementary Material, ‘Methods’ section)**.

## Results

To test whether electrical stimulation of pulvinar subregions selectively engages occipital, lateral temporal, and parietal pathways, we applied SPES to sEEG contacts placed in the PuL, vPuM, and dPuM pulvinar subregions (**Figure 1a)**. A smoothed group map of stimulation to the PuL, vPuM and dPuM shows the effective connectivity of these regions to the cortex. The effective connectivity shows that PuL has strong influence on occipital brain regions, vPuM has strong influence on lateral temporal areas and less strong influence on occipital areas whereas dPuM has strong influence on relatively more anterior lateral temporal areas and parietal areas. These maps indicate that stimulation of pulvinar subregions may influence distinct cortical areas.

**Figure 1.**
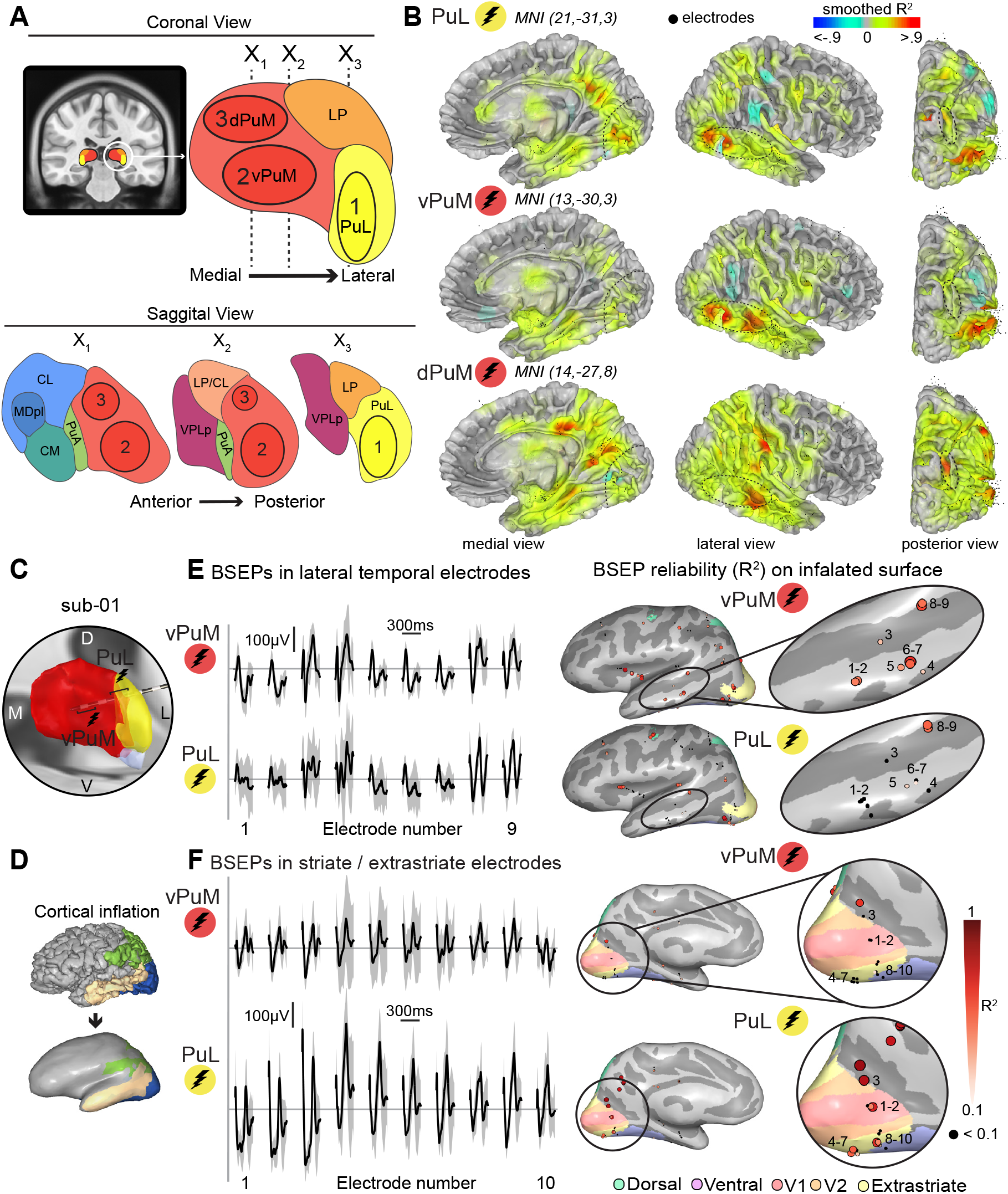
Stimulation of pulvinar subregions shows preferential influence on distinct cortical areas. **A)** Pulvinar schematic showing three pulvinar subregions: lateral pulvinar (PuL), ventromedial pulvinar (vPuM) and dorsomedial pulvinar (dPuM). LP=lateral posterior nucleus, MDpl=mediodorsal nucleus (paralamellar), CL=central lateral nucleus, CM=centromedian nucleus, VPLp=ventral posterior lateral nucleus (anterior division), PuA=anterior pulvinar. B) Rendering of smoothed average brain stimulation evoked potentials (BSEP) consistency (R^2^) across subjects shows that PuL stimulation has a reliable influence on striate and extrastriate (visual) areas, vPuM has a reliable influence on lateral temporal cortex and dPuM has a reliable influence on parietal cortex. **C)** Electrode trajectory for sub-01 D) Cortical inflation schematic illustrating how inflated brain surfaces are created. **E)** Stimulation of vPuM and PuL have different brain stimulation evoked potentials (BSEPs) in sub-01 when measuring in the lateral temporal cortex. Black lines indicate mean BSEP responses across trials, and gray lines are bootstrapped 95% confidence intervals in nine different measurement electrodes. Inflated brain surfaces show pulvinar-cortical effective connectivity on electrodes measuring in the lateral temporal cortex when stimulating vPuM and PuL. There are reliable and larger responses only when stimulating vPuM. **F)** vPuM and PuL stimulation in the same subject but now measuring pulvinar-cortical effective connectivity in striate and extrastriate electrodes. Visual electrodes have larger BSEP amplitude and pulvinar-cortical effective connectivity in ten electrodes when receiving stimulation from PuL.

The selectivity between PuL and vPuM was also observed in a single subject. In sub-01, electrodes were placed in the vPuM and PuL (**Figure 1b**). The nine electrodes measuring from the lateral temporal cortex show robust BSEPs when stimulating vPuM, with amplitudes of >100μV (**Figure 1d top**). Stimulation of PuL, however, shows weaker evoked responses of <100μV in amplitude in the same recording electrodes measuring from the lateral temporal cortex (**Figure 1d bottom**). As a measure of the strength of the pulvinar-cortical effective connectivity, we calculate the BSEP consistency (R^2^), which is high if stimulation elicits a similar BSEP response across trials. Rendering R^2^ on the inflated brain surface (**Figure 1c**) shows that the lateral temporal cortex exhibits consistent BSEPs in response to vPuM stimulation and less consistent BSEPs in response to PuL stimulation (**Figure 1d right**).

Within the same subject, an additional ten electrodes were used to measure BSEPs from striate and extrastriate areas. Stimulation of vPuM elicits weak BSEPs, typically <150μV in amplitude within these early visual cortical areas (**Figure 1e, top)**. However, stimulation of PuL elicits robust BSEPs, often exceeding 250μV in amplitude (**Figure 1e, bottom)**. Rendering BSEP consistency (R^2^) on an inflated brain surface shows that vPuM stimulation results in a weak outgoing influence on early visual cortex, while PuL stimulation results in a strong outgoing influence on the same visual measurement electrodes (**Figure 1e, right)**.

### Proximity to medial pulvinar increases strength of pulvinar-cortical influence to lateral temporal cortex

Previous pulvinar studies have suggested that the vPuM has robust connections to the lateral temporal cortex.^15-16^ To causally evaluate this claim and further determine if there is a subregion within the pulvinar (**Figure 2a)** that is effectively connected to the lateral temporal cortex, we parameterize BSEPs and plot the pulvinar-cortical effective connectivity strength from three example lateral temporal electrodes in a bar chart (**Figure 2b)**. The BSEP consistency (R^2^) in the lateral temporal cortex is the largest in response to vPuM stimulation. As stimulation transitions from vPuM to PuL, the observed pulvinar-cortical effective connectivity strength diminishes, typically remaining < 0.3. When rendering BSEP influence on an inflated brain surface and comparing the most medial and lateral stimulation sites (**Figure 2b, right)**, our data shows that the most ventral-medial stimulation sites drive the strongest pulvinar-cortical influence to the lateral temporal cortex. This is further demonstrated in an exemplary subject (**Figure 2c**) where the medial most pulvinar stimulation strongly influences lateral temporal regions. This relationship is also evident within the vPuM, where the distance from the medial aspect appears to mediate the response influence, which recedes posteriorly and diminishes as stimulation approaches the lateral pulvinar.

**Figure 2.**
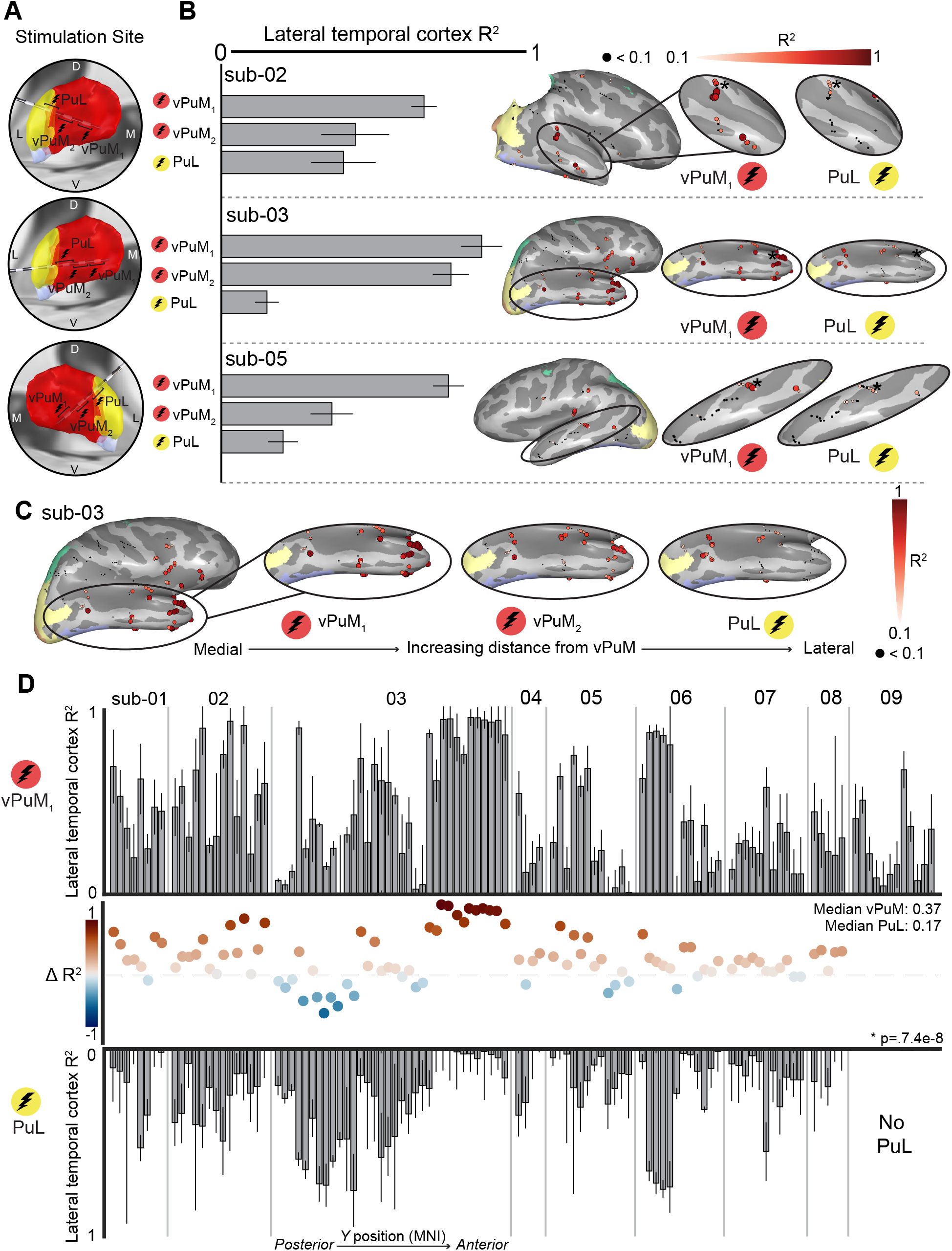
Proximity to ventromedial pulvinar increases pulvinar-cortical effective connectivity strength to the lateral temporal cortex. **A)** Stimulation sites for three subjects receiving pulvinar stimulation. **B)** Measurement of pulvinar-cortical effective connectivity strength (R^2^) in an example lateral temporal cortex measurement electrode (asterisk). Ventromedial pulvinar (vPuM) stimulation delivers higher R^2^ than lateral pulvinar (PuL) stimulation. Inflated surfaces show pulvinar-cortical influence for all cortical electrodes in three subjects. Each dot represents an electrode, and size and color represent R^2^. We compare the most medial pulvinar stimulation (red) with the most lateral pulvinar stimulation (yellow). **C)** Pulvinar-cortical influence across all stimulation sites in sub-03 **D)** Measurement of pulvinar-cortical effective connectivity strength across all lateral temporal measurement electrodes. The top bar chart shows the most medial pulvinar stimulation (red), and the bottom is the most lateral pulvinar stimulation. Dots along the middle show the difference (vPuM-PuL) between pulvinar-cortical effective connectivity strength. Red indicates that most medial pulvinar stimulation effective connectivity is stronger to the lateral temporal cortex than most lateral pulvinar stimulation; blue indicates the opposite.

To show that this is a consistent effect, we then visualize the parameterized BSEP consistency from all nine subjects with lateral temporal electrodes and compare the most medial (**Figure 2d, top)** and most lateral (**Figure 2d, bottom)** pulvinar stimulation sites. Stimulation in the vPuM elicits significantly stronger influence on the lateral temporal cortex compared to stimulation of the PuL (Wilcoxon signed-rank test, p=7.4e-8, median vPuM R^2^ = 0.37, median PuL R^2^ =0.17). We also take the difference between parameterized BSEP effective connectivity in the most medial and most lateral (vPuM-PuL) stimulation sites (**Figure 2d, middle)**. This allows us to visually determine that in 76% (76/100) of lateral temporal cortex electrodes, BSEPs had a more substantial influence on the lateral temporal cortex when stimulating the most medial vPuM electrode pair than the most lateral PuL stimulation pair.

### Proximity to lateral pulvinar PuL increases strength of pulvinar-cortical influence to striate and extrastriate areas

The lateral pulvinar is known to be connected to striate and extrastriate areas, with particular visual connectivity noted in the inferior portion of the lateral pulvinar.^12-15^ To causally test whether direct PuL stimulation influences these areas in the human brain, we parameterized BSEPs after stimulation of different pulvinar sites (**Figure 3a)**. A bar graph in two examples of striate/extrastriate visual electrodes shows that across vPuM and PuL stimulation, pulvinar-cortical effective connectivity strength is largest when stimulating the closest electrode to PuL (**Figure 3b)**. In example subject 2, a comparison of vPuM and PuL stimulation shows that BSEP consistency is higher in visual areas when stimulating the lateral pulvinar (**Figure 3c)**. This relationship is similarly shown for exemplary subject 3 (**Figure 3d)**. Interestingly, lateral pulvinar stimulation did not selectively influence subsets of visual areas but elicited responses across many electrodes in striate and extrastriate regions.

**Figure 3.**
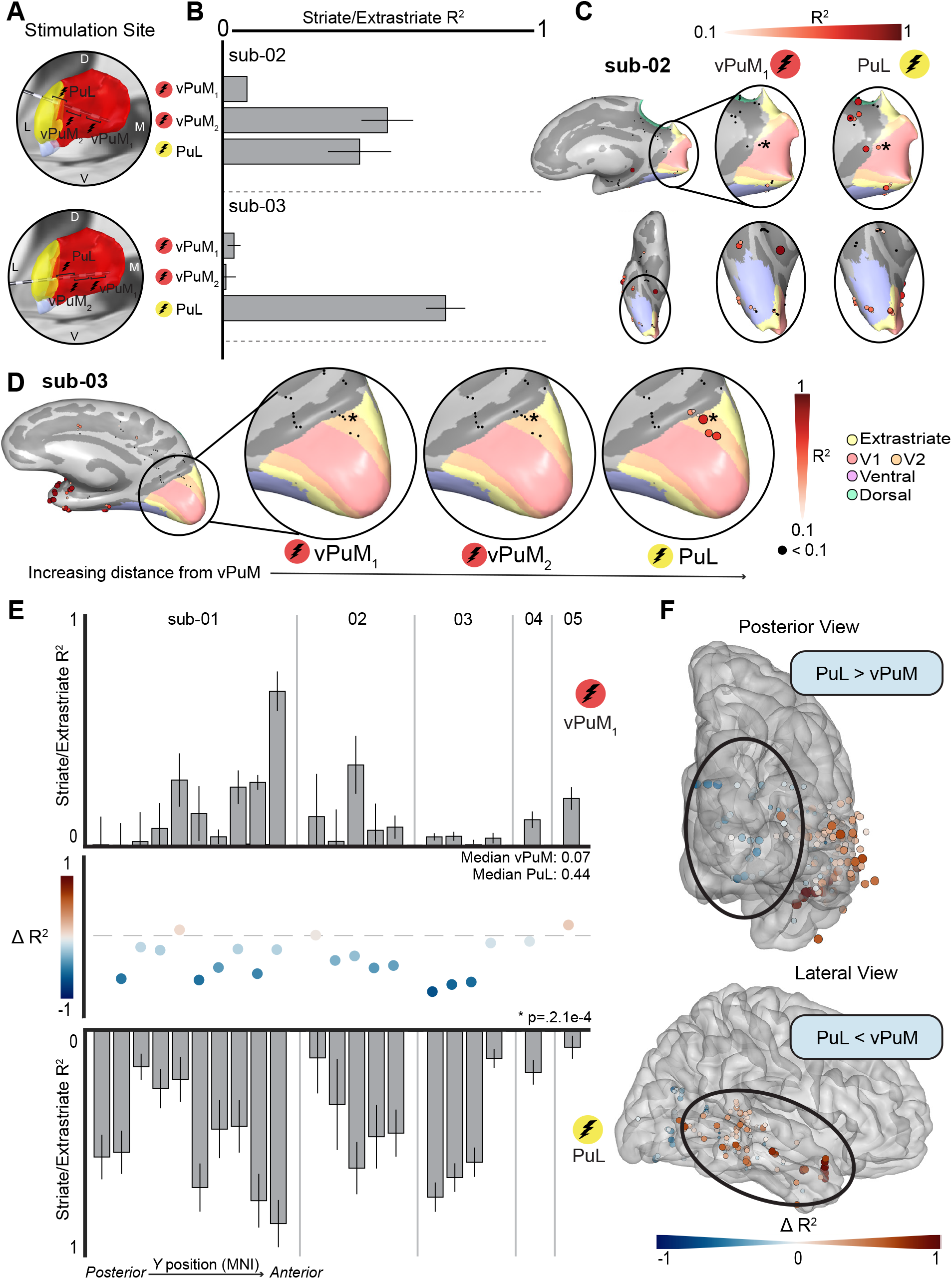
Proximity to lateral pulvinar increases pulvinar-cortical effective connectivity strength to visual areas. **A)** Stimulation sites for two subjects receiving pulvinar stimulation. **B)** Pulvinar-cortical effective connectivity strength (R^2^) of one example striate/extrastriate measurement electrode (asterisk). Most medial (vPuM) stimulation is less than most lateral (PuL) stimulation. **C-D)** Example sub-02/sub-03 comparing vPuM stimulation and PuL stimulation regarding pulvinar-cortical effective connectivity strength. **E)** Measurement of pulvinar-cortical effective connectivity strength across all striate/extrastriate measurement electrodes. The top bar chart shows most medial pulvinar stimulation (red), the bottom is most lateral pulvinar stimulation (yellow). Dots along the middle show the difference (vPuM-PuL) between pulvinar-cortical effective connectivity strength. Blue indicates most medial pulvinar stimulation influence is weaker to the lateral temporal cortex than most lateral pulvinar stimulation; red indicates the opposite. **F)** MNI warp of all electrodes across subjects showing the difference between most medial and most lateral pulvinar stimulation (blue/red dots from Fig2D and 3E). The lateral temporal cortex shows stronger pulvinar-cortical influence from vPuM, whereas posterior visual electrodes receive stronger influence from PuL.

Next, we visualized the parameterized BSEP consistency from all five subjects with striate and extrastriate electrodes to compare BSEP responses to the most medial (**Figure 3e, top)** and most lateral (**Figure 3d, bottom)** pulvinar stimulation sites. Stimulation in the vPuM elicited significantly less influential responses in striate and extrastriate electrodes compared to stimulation of the PuL (Wilcoxon signed-rank test, p=2.1e-4, median PuL=0.44, median vPuM=0.07). We also investigate the difference between parameterized BSEP effective connectivity in response to the most medial and most lateral stimulation (vPuM-PuL) sites (**Figure 3e, middle)**. This allows us to visually determine that in 86% (18/21) of striate and extrastriate electrodes, BSEPs had a smaller influence when stimulating the most medial vPuM electrode pair than the most lateral PuL stimulation pair. Of note, in subjects 4 and 5, these effects were less pronounced compared to subjects 1, 2, and 3. This may be because in subjects 4 and 5, electrodes were placed more dorsal in the PuL compared to the more inferior locations in subjects 1, 2, and 3 (Supplemental Figure 3). Stimulation of the inferior portion of the lateral pulvinar stimulation thus strongly influenced striate and extrastriate areas.

Illustrating the difference in pulvinar-cortical effective connectivity (vPuM-PuL) revealed two clusters (**Figure 3f)** in MNI-152 space when visualizing all responsive cortical electrodes in striate/extrastriate and lateral temporal cortex electrodes. PuL stimulation exhibits enhanced effective connectivity strength with electrodes in the striate and extrastriate cortex, while vPuM stimulation exhibits effective connectivity strength with lateral temporal cortex.

### The dorsomedial pulvinar influences parietal regions

Previous anatomical studies have indicated that the dorsal pulvinar is most strongly connected to the intraparietal sulcus (IPS) and several frontal brain regions like the insulo-opercular cortices.^15-16^ To evaluate the selectivity of the dPuM, we stimulated dPuM in three subjects who had electrodes placed in the dPuM as well as in the IPS, striate, extrastriate, and lateral temporal cortex. Stimulation of these dPuM sites elicited significant responses in IPS, with amplitudes of > 50 μV, as well as in insulo-opercular electrodes (**Figure 4a, b, top** for subjects 10 and 11, subject 12 shown in Figure S4c). However, responses in the striate, extrastriate, and lateral temporal cortex were much more limited, with typical amplitudes of < 25 μV (**Figure 4a, b, bottom)**. The pulvinar-cortical effective connectivity visualized on the inflated surfaces shows reliable responses tend to aggregate in the parietal dorsal stream (IPS1-5) and in the insulo-opercular cortex, with limited dPuM influence in striate/extrastriate or lateral temporal cortex. As these subjects did not have electrodes in the vPuM or PuL, we could not directly compare the inputs from different pulvinar subregions. However, across four subjects with varying dPuM-vPuM trajectories, stimulation of the dPuM engaged parietal areas, whereas stimulation of the vPuM shifted towards lateral temporal engagement (**Figure 4c**). These data indicate that dorsal pulvinar stimulation may have a relatively selective influence on parietal areas without strong striate/extrastriate and lateral temporal cortex engagement.

**Figure 4.**
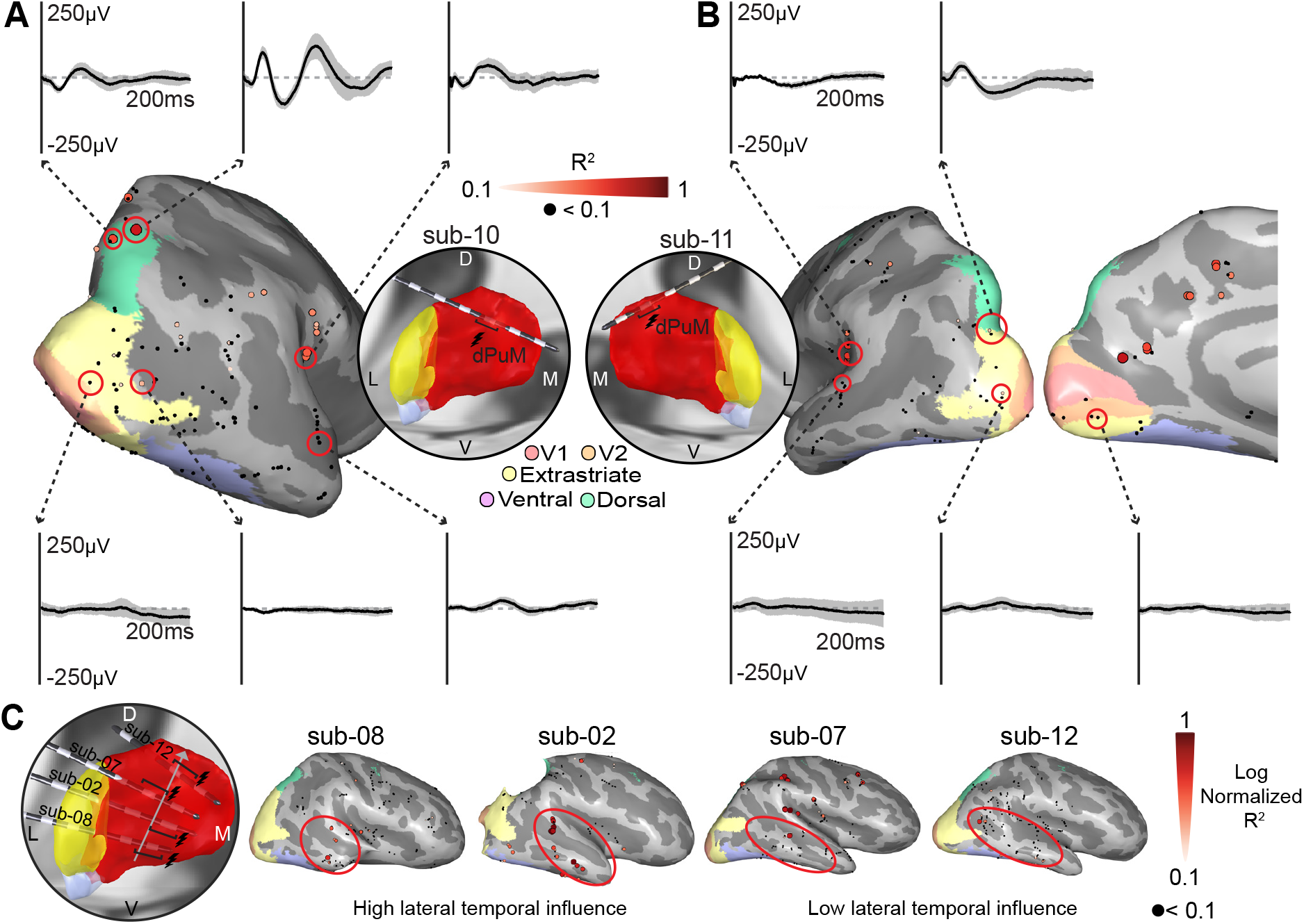
The dorsomedial pulvinar (dPuM) influences parietal and insulo-opercular cortices. **A)** Example brain stimulation evoked potentials (BSEPs) from sub-10. There are reliable BSEPs from the dorsomedial pulvinar to IPS cortices and insulo-opercular areas, with large effective connectivity strength (R^2^). Striate/extrastriate and lateral temporal cortex BSEPs are limited. Black lines represent average BSEPs, and shaded gray lines represent a bootstrapped 95% confidence interval. **B)** Example BSEPs from sub-11. Similarly to sub-10, there is reliable parietal pulvinar-cortical effective connectivity, with limited lateral temporal cortex or striate/extrastriate influence. **C)** Log normalized effective connectivity strength of multiple subjects. Stimulation of ventral locations has high lateral temporal influence, which is lost as we move towards the dorsal pulvinar.

## Discussion

### Pulvinar stimulation drives select zones of effective connectivity

In this study, we investigated whether stimulation of pulvinar subregions selectively influences lateral temporal, striate/extrastriate, and parietal cortical areas. By stimulating the pulvinar with SPES in twelve subjects, we found that pulvinar effective connectivity follows several motifs. Stimulation of PuL evoked reliable BSEPs in striate and extrastriate electrodes preferentially over vPuM stimulation. Stimulation of vPuM evoked strong BSEPs in the lateral temporal cortex, decreasing as stimulation progressed laterally. Stimulation of the dPuM, on the other hand, evoked parietal responses without robust early visual or lateral temporal responses. While these gradients were not always selective, stimulation of different pulvinar areas showed strong response preferences for select cortical areas.

### Pulvinar stimulation zones of effective connectivity follow anatomical pathways

The measured pulvinar effective connectivity corresponds well with previous anatomic delineations. Tracer studies in macaques found that the lateral and inferior pulvinar most strongly project to striate and extrastriate areas.^12-13^ The medial pulvinar, however, projects to temporal, parietal, frontal, orbital, and cingulate cortices.^13-14, 21-22^ Furthermore, human studies leveraging diffusion imaging and fMRI have found similar structural and functional connectivity between pulvinar nuclei and select cortical regions.^15-16,23^ These studies proposed two topographical gradients for pulvinar connectivity. ^12-14^ First, PuL has predominantly early visual connectivity, while PuM is connected to higher-order (temporal, parietal) cortices. Second, the ventral pulvinar has predominantly visual and temporal outputs, while the dorsal pulvinar has stronger parietal outputs. Our stimulation data mirrors these two axes. These findings can provide guidance when implanting pulvinar electrodes intended to influence select cortical regions.

### Pulvinar anatomy has to be considered in clinical pulvinar targeting

SPES provides causal insight into better understanding the extent to which pulvinar stimulation influences select anatomical pathways. Initially, pulvinar stimulation was proposed for temporal neocortical epilepsy; recently, studies have used pulvinar stimulation more generally for the posterior quadrant and other epilepsies.^6, 8-10^ Previous studies have indicated that thalamic DBS in epilepsy may be most effective when stimulation is applied to thalamic structures that project to seizure onset regions.^11,24^ Our data suggest that dPuM stimulation, for example, may not be very effective in influencing neural activity in the posterior quadrant or lateral temporal areas, given the relatively limited effective connectivity between the dPuM and these regions. This is especially important to consider given the vast heterogeneity of clinical symptoms for patients with drug-resistant epilepsy in the posterior quadrant or lateral temporal cortex.

### Directionality of pulvinar connectivity

Pulvinar to cortical connectivity is often described as highly bi-directional.^13,25-26^ However, previous literature suggests that connectivity is not always bi-directional.^16^ For example, studies with medial pulvinar stimulation showed that hippocampal electrodes only responded 16% of the time.^16^ Our supplemental data (**Supplemental Figure 5)** shows that hippocampus BSEPs from pulvinar are similarly found in very few sites and subjects (17% electrode responses across subjects). The BSEPs are relatively muted except for one patient whose electrodes were placed close to the fornix (Supplemental Figure 5b). These sparse pulvinar-hippocampal outputs are opposite to strong hippocampal-pulvinar outputs; evidence shows that hippocampal stimulation or seizures commonly propagate to the medial pulvinar.^16,27^ The fact that seizures spread to the pulvinar, therefore, does not necessarily make the pulvinar an ideal hippocampal DBS target, as pulvinar stimulation may have little to no effect on a mesial temporal seizure onset zone. Because of this discrepancy, it remains essential to consider pulvinar anatomy and use electrical stimulation to test whether seizure onset regions can be effectively driven with pulvinar stimulation.

### Limitations

Stereotactic EEG provides sparse spatial coverage of the brain, and this study can only evaluate cortical areas that were well sampled; we were unable to derive results on how pulvinar stimulation influenced cortical areas that were under-sampled or not sampled. Our results, therefore, focus on the differences in evoked responses in the same cortical electrodes when stimulating different pulvinar nuclei. This approach is restricted in that it does not allow us to prove a lack of connection between unsampled regions. However, it did allow us to compare the relative influence of inputs from different pulvinar areas.

Our approach did not make any assumptions about BSEP waveform shape or latency, and it is possible that some evoked potentials were generated by indirect connections. Furthermore, electrical stimulation can result in orthodromic as well as antidromic activation.^28^ While our parameterization algorithm (CRP) performs a direct quantification of similarities between different trials, we did not assess the consistency or similarity of BSEP shapes across subjects. More detailed waveform analysis may provide an increased understanding of the local neuronal population, or the architecture of the signal being sent from pulvinar subregions to cortical areas. Consistent waveform differences may inform whether BSEPs contained indirect or antidromic activation.

Lastly, although these short-latency electrophysiological biomarkers demonstrate causal stimulation and measurement in neuronal populations, it remains unclear whether these biomarkers are useful for predicting long-term clinical outcomes of neuromodulation devices in the targeted regions.^29^

## Conclusion

We show that brain stimulation evoked potentials can be used to map topographically organized effective connectivity fields of the pulvinar nucleus. This pulvinar subfield mapping may be used to tailor brain stimulation to engage intended lateral temporal and posterior quadrant networks, which is relevant for the potential future of individualized neuromodulation for epilepsy. Future work should assess long-term therapeutic outcomes and compare the impact of targeted vs conventional pulvinar DBS.

## Code and data accessibility

All analyses were performed in MATLAB 2023b. Code will be available on our GitHub (https://github.com/MultimodalNeuroimagingLab) page in addition to de-identified data to reproduce the main figures on the Open Science Framework (that will be made available with manuscript acceptance).

## Acknowledgements

The authors would like to thank the patients who participated in this study, Karla Crocket, Cindy Nelson and other staff and colleagues who assisted at Saint Marys Hospital and Barnes-Jewish Hospital. We thank Zeeshan Qadir for valuable discussions. We also thank Michael Jensen and Gabriela Ojeda Valencia for their help with data curation.

