## Supplemental Table 1, Supplemental Table 2 for "Human pulvinar stimulation engages select cortical pathways in epilepsy"

### Methods

#### Electrode localization

T1-weighted (T1w) images were aligned and resliced along their anterior-posterior commissure (ACPC) axis, which was subsequently segmented using FreeSurfer's cortical reconstruction and manually inspected.[1] The postoperative CT was coregistered onto the preoperative ACPC T1w image, and electrode positions were labeled according to the FreeSurfer Destrieux Atlas to calculate hippocampal response rates.[2] Electrodes were additionally visualized using LeadDBS 3.0, and pulvinar electrodes were identified by using the multiarchitectonic and stereotactic Krauth/Morel atlas; included sites were electrodes within the medial and lateral pulvinar.[3-5] The anterior and inferior pulvinar were not sampled as part of this study due to clinical electrode placement. Further visual-cortex specific atlases were applied using the neuropythy library.[6-8]

#### Selection of significant brain stimulation evoked potential

Significant brain stimulation evoked potentials were determined after parameterizing the responses from differing pulvinar stimulation site. We employed three main criteria from the Canonical Response Parameterization (CRP) algorithm.[9]

1) Median coefficient of determination greater than 0.1

2) Mean signal-to-noise ratio (ratio of projection weight to residual) greater than 0.5

3) P-value less than 0.05 (t-test of cross-projection magnitude at significant response time vs zero)

After significant brain stimulation evoked potentials were identified, they were subject to a manual review (J.A.B). This ensured that no trials with reliable artifacts or synchronous epileptiform events that were biasing the parameterization algorithm towards a significant response when no such response was present. These channels were discarded and not used for analysis.

#### Selection of cortical electrodes

Intracortical electrodes within 6mm of the surface were projected to the nearest cortical vertex. Cortical striate (V1-V2), extrastriate (V3a/b, hV4, TO1-2, LO1-2, IPS0), and parietal (IPS1-5) electrodes were included based on placement within Wang, Rosenke, and Benson atlases regions of interest. A file describing selected electrodes can be found: /derivatives/stats/pulvinar_stats.xlsx

### Discussion

#### Stimulation parameters influence pulvinar-cortical effective connectivity strength

It is important to understand whether stimulation parameters influence pulvinar-cortical effective connectivity fields. Different groups use varying single-pulse electrical stimulation parameters, which can often drive differing responses. We describe two types of effects that are important to consider when changing stimulation parameters to map pulvinar effective connectivity. First, we consider changes in the influence of intended targets and second, we consider the fact that a larger current increases the volume of tissue activation and therefore may increase off-target effects.

First, we show BSEPs for one subject who received a combination of stimulation parameters **(Supplemental Figure, 4a-b)**. When applying a small amplitude 1mA ventromedial pulvinar (vPuM) pulse with a width of 100µs or 200µs, BSEPs in the lateral temporal cortex typically exhibit response amplitudes < 150µV (**Supplemental Figure, 4c**). However, as the pulse amplitude increases up to 6mA, BSEP amplitude in the lateral temporal cortex also increases, exhibiting response amplitudes greater than 200µV and maximal 800µV. When stimulating in the vPuM while increasing the charge density, pulvinar-cortical influence increases in the lateral temporal cortex. Notably, stimulation of 3mA-200µs and 6mA-100µs has an equal charge density but elicits different responses. While 3mA-200µs has broad activations, 6mA-100µs is strongly selective for particular electrodes. **(Supplemental Figure, 4d).**

Second, across the three subjects we stimulated with variable pulse widths, effects differed **(Supplemental Figure, 4e-f).** Larger pulse width (200µs) caused more variable effects in 2 of the 3 subjects **(Supplemental Figure, 4f)** where pulvinar effective connectivity fields with increased current varied to those mapped at 100µs. In these two subjects, the electrodes were positioned closer to the extremity of the pulvinar compared to the subject where this was not the case. The higher stimulation current may be engaging unintended structures. Therefore, when trying to understand whether pulvinar subregions influence specific anatomical areas, it remains important to consider stimulation parameters when analyzing the strength and spread of the cortical influence.

#### Supplementary Tables

| Subject | Age | Sex | #Electrodes/ #Leads/Hemisphere | Pulvinar Electrodes | Lesions/ Seizure Onset Zones(s) |
|---|---|---|---|---|---|
| sub-01 | 36 | M | 234/15/B | 4 | **Lesion**: Occipital encephalomalacia<br><br>**SOZ**: Occipital encephalomalacia, hippocampus |
| sub-02 | 15 | M | 217/15/B | 4 | **Lesion**: Posterior paramedian<br><br>**SOZ**: Inferior occipito-temporal (fusiform), posterior cingulate |
| sub-03 | 39 | M | 231/19/R | 4 | **Lesion:** Posterior temporal/supramarginal and anterior inferiorparietal.<br><br>**SOZ:** Anterior perilesional |
| sub-04 | 23 | F | 221/15/B | 4 | **Lesion**: n/a<br><br>**SOZ**: Mesial temporal, temporal neocortex |
| sub-05 | 38 | M | 181/13/R | 4 | **Lesion**: Inferior temporal focal cortical dysplasia<br><br>**SOZ**: Posterior lesion, hippocampus |
| sub-06 | 18 | M | 222/14/L | 4 | **Lesion**: Inferior temporal encephalomalacia<br><br>**SOZ**: Inferior temporal |
| sub-07 | 27 | M | 243/16/B | 3 | **Lesion**: n/a<br><br>**SOZ**: Fusiform, occipito-temporal sulcus |
| sub-08 | 21 | M | 231/14/R | 5 | **Lesion**: Temporal lobectomy, amygdalohippocam-pectomy<br><br>**SOZ**: Insula |
| sub-09 | 16 | M | 256/18/B | 3 | **Lesion**: n/a |

| | | | | | |
|---|---|---|---|---|---|
| | | | | | **SOZ**: Posterior operculum, temporal neocortical, mesial temporal |
| sub-10 | 41 | F | 168/12/R | 2 | **Lesion**: Prior dysembryoplastic neuroepithelial tumor resection (occipito-mesial temporal)<br><br>**SOZ**: Resection cavity |
| sub-11 | 24 | M | 192/13/B | 3 | **Lesion**: Medial occipital encephalomalacia,<br><br>**SOZ**: Middle temporal gyrus, encephalomalacia |
| sub-12 | 29 | F | 231/16/B | 3 | **Lesion**: Posterior superiotempral gyrus<br><br>**SOZ**: Anterior perilesional |

**Supplementary Table 1. Subject information.** Table details age, sex, number of contacts, leads, laterality, number of pulvinar electrodes, lesions, and primary seizure onset zones. M=male, F=female, R=right, L=left, B=bilateral.

| Subject | Amplitude (mA) | Pulse Width (µs) | Sampling Rate (Hz) | Number of trials per pulvinar electrode | Amplifier | Cortical Stimulator |
|---|---|---|---|---|---|---|
| sub-01 | 6 | 200 | 2048 | 12-24 | Natus Quantum | Natus Nicolet |
| sub-02 | 6 | 100 | 4800 | 12 | g.tec HIAMP | g.tec ESTIMPRO |
| sub-03 | 1, 3, 6 | 100, 200 | 2000 | 60 | Nihon Kohden JE120A | Nihon Kohden MS-120BK-EEG |
| sub-04 | 6 | 100 | 4800 | 24 | g.tec HIAMP | g.tec ESTIMPRO |
| sub-05 | 6 | 100 | 4800 | 12 | g.tec HIAMP | g.tec ESTIMPRO |
| sub-06 | 6 | 100, 200 | 4800 | 36, 12, respectively | g.tec HIAMP | g.tec ESTIMPRO |
| sub-07 | 6 | 100, 200 | 4800 | 36 | g.tec HIAMP | g.tec ESTIMPRO |
| sub-08 | 6 | 100 | 4800 | 12 | g.tec HIAMP | g.tec ESTIMPRO |
| sub-09 | 6 | 100 | 4800 | 12 | g.tec HIAMP | g.tec ESTIMPRO |
| sub-10 | 6 | 100 | 4800 | 12 | g.tec HIAMP | g.tec ESTIMPRO |
| sub-11 | 6 | 100 | 4800 | 12 | g.tec HIAMP | g.tec ESTIMPRO |
| sub-12 | 1, 3, 6 | 100, 200 | 2000 | 60 | Nihon Kohden JE120A | Nihon Kohden MS-120BK-EEG |

**Supplementary Table 1. Stimulation conditions.** The table details amplitude (mA), pulse width (µs), sampling rate (Hz), number of stimulation trials delivered per pulvinar electrode, amplifier, and cortical stimulator per subject. Multiple stimulation amplitudes or pulse widths for each subject are a combination (ex: sub-03 had 60 trials of 1mA-100µs, 3mA-100µs, 6mA-100µs, 1mA-200µs, 3mA-200µs, and 6mA-200µs). Sub-01 had between 12-24 trials of pulvinar stimulation electrodes. Sub-06 had 36 trials for 6mA-100µs and 12 for 6mA-200µs.

#### Supplementary Figures

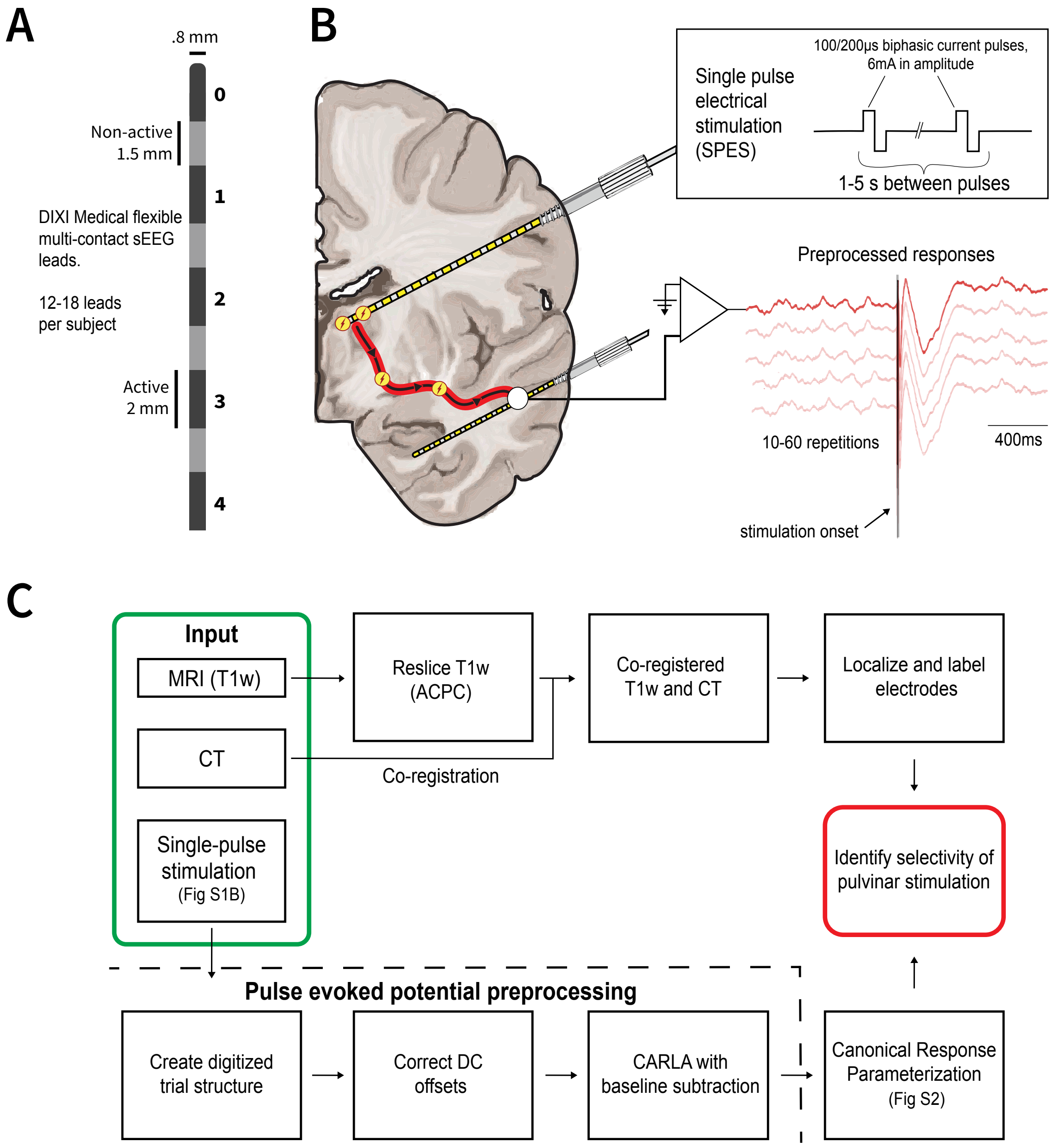


**Supplemental Figure 1 Electrode diagram and schematic of single pulse electrical stimulation protocol. A)** Diagram of DIXI multi-contact stereotactic EEG (sEEG) lead. Contacts are 2mm in length with a 1.5mm gap between contacts; the entire lead has a diameter of .8mm. Numbers are placed on the midpoint between contacts and refer to the bipolar contact pair. **B)** Schematic of single pulse electrical stimulation (SPES) illustrates thalamocortical stimulation from a bipolar contact pair. Stimulation parameters are many (**See Supplemental Table 2)** and occur every 1-5s between pulses. They are jittered within time, and 12-60 stimulation repetitions are performed. **C)** Schematic illustrating preprocessing and analysis pipeline.

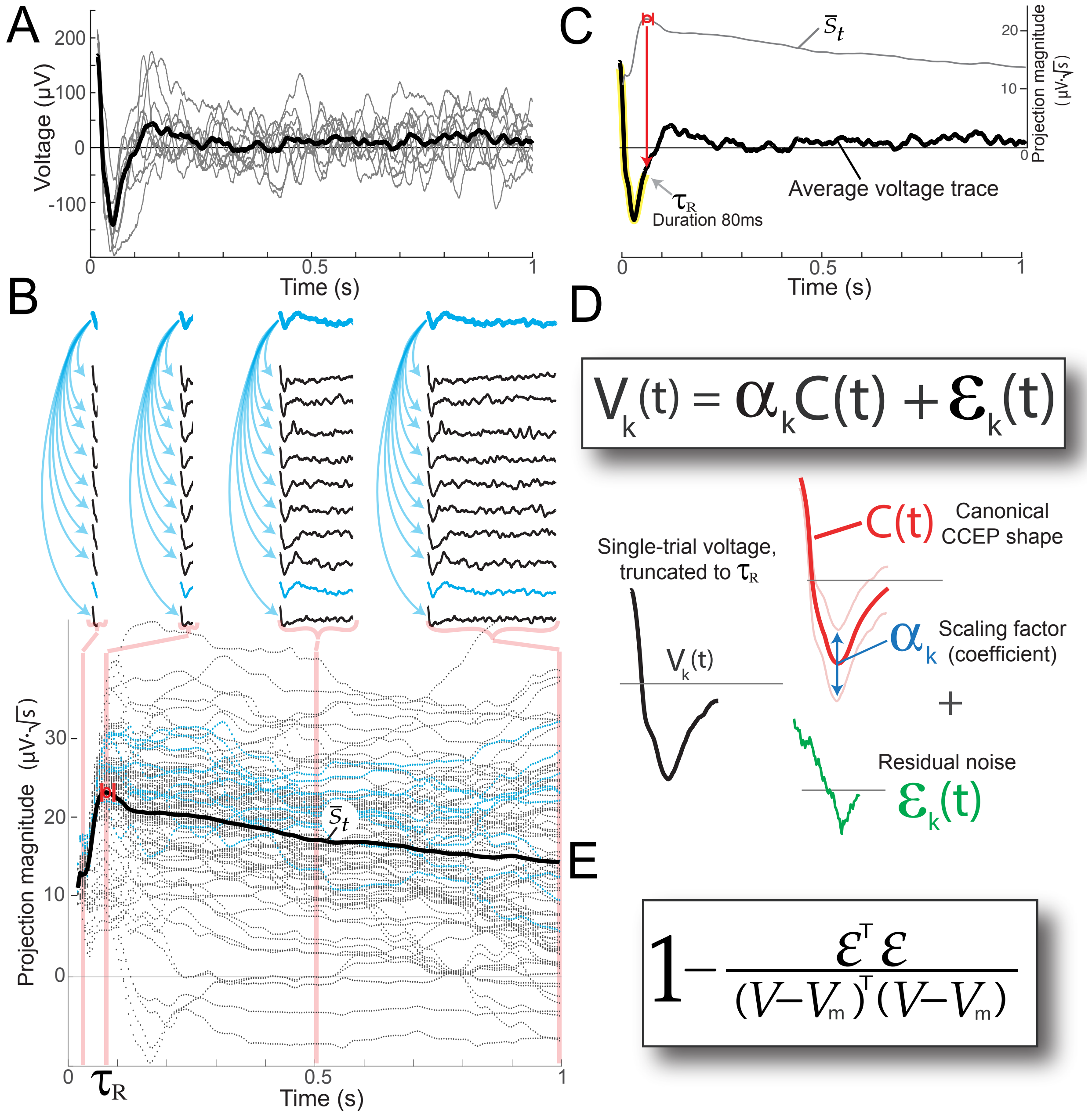


**Supplemental Figure 2 Illustration of CRP method. A)** Individual trials are recorded, between 10-60 repetitions, and an average trace is computed. **B)** A matrix is calculated where each M unit-normalized trial is projected against all other trials for N time points with self-projections removed. **C)** The matrix is sorted into a set that provides a distribution of cross-projection magnitudes, while the average projection magnitudes summarize the interaction from stimulation to response. The maximum of this distribution in time is used to identify the significant response time ($\tau_R$), highlighted in yellow. A linear kernel PCA can be applied to the individual trials truncated to the $\tau_R$ which captures the canonical CCEP C(t). **D)** One trial $V_k(t)$ is equivalent to the canonical CCEP C(t) multiplied by a scaling coefficient ($\alpha_k$) added to residual noise $\varepsilon_k(t)$. **E)** The residual noise, or “residuals” can be used to calculate the coefficient of determination ($R^2$) of the brain stimulation evoked potential as a measure of pulvinar-cortical effective connectivity. For more details refer to the original Miller *et al*.[9] or Ojeda Valencia *et al*.[10] for use case.

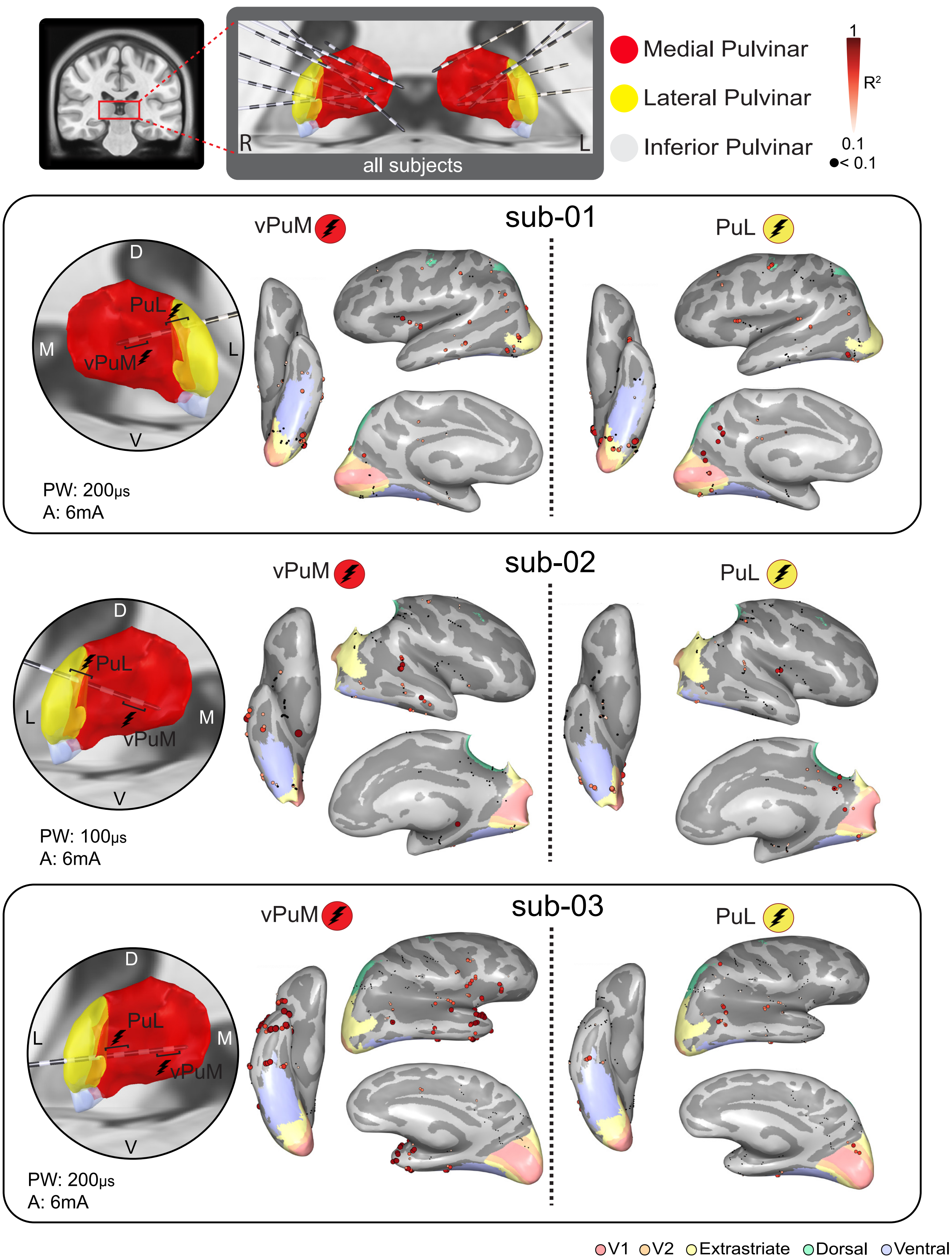

Medial Pulvinar
Lateral Pulvinar
Inferior Pulvinar
1
$R^2$
0.1
< 0.1
R
L
all subjects
sub-01
vPuM
PuL
D
M
L
V
PW: 200μs
A: 6mA
sub-02
vPuM
PuL
D
L
M
V
PW: 100μs
A: 6mA
sub-03
vPuM
PuL
D
L
M
V
PW: 200μs
A: 6mA
V1
V2
Extrastriate
Dorsal
Ventral

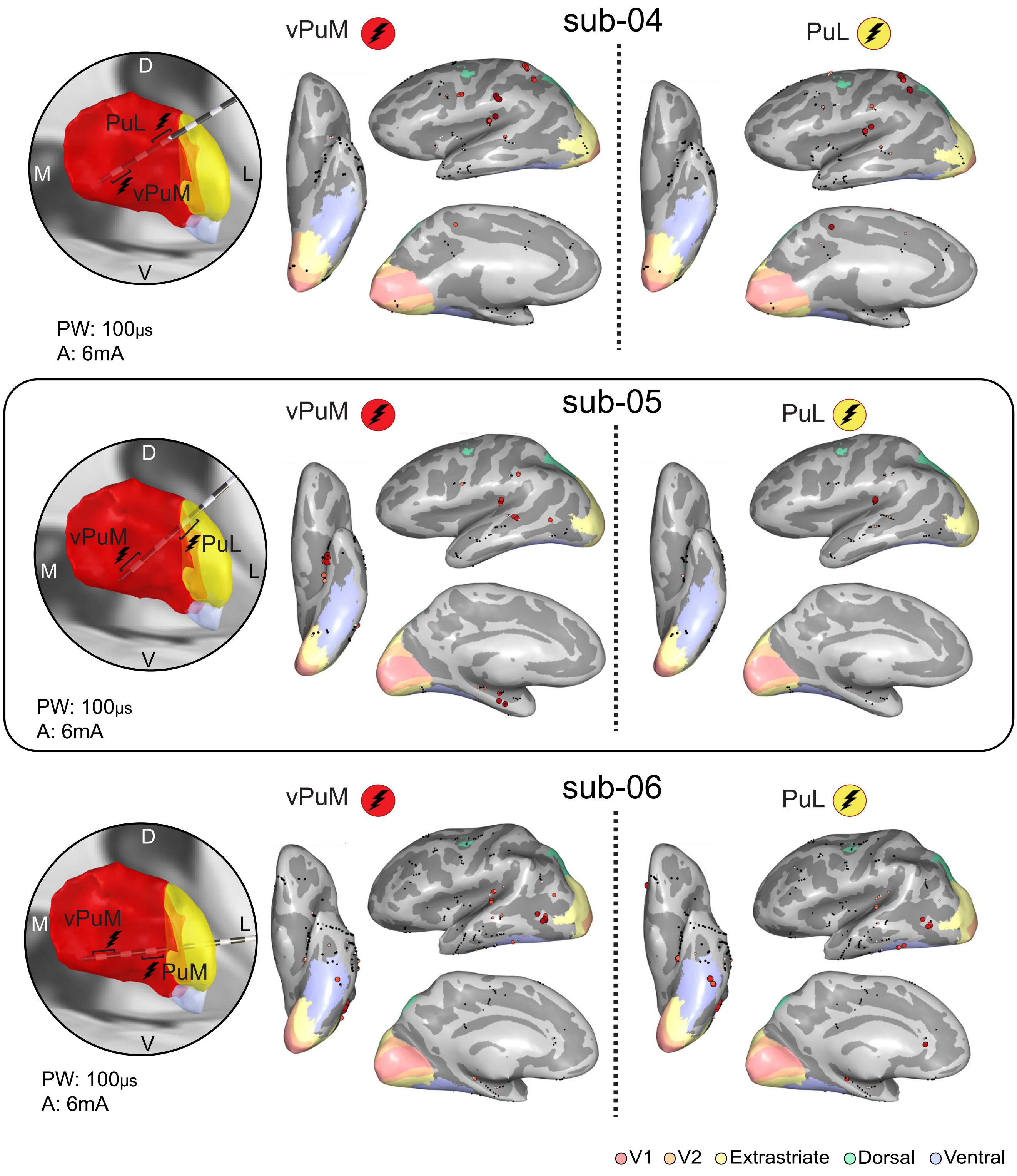
sub-04
vPuM
PuL
D
M
L
V
PuL
vPuM
PW: 100μs
A: 6mA
sub-05
vPuM
PuL
D
M
L
V
vPuM
PuL
PW: 100μs
A: 6mA
sub-06
vPuM
PuL
D
M
L
V
vPuM
PuM
PW: 100μs
A: 6mA
V1
V2
Extrastriate
Dorsal
Ventral

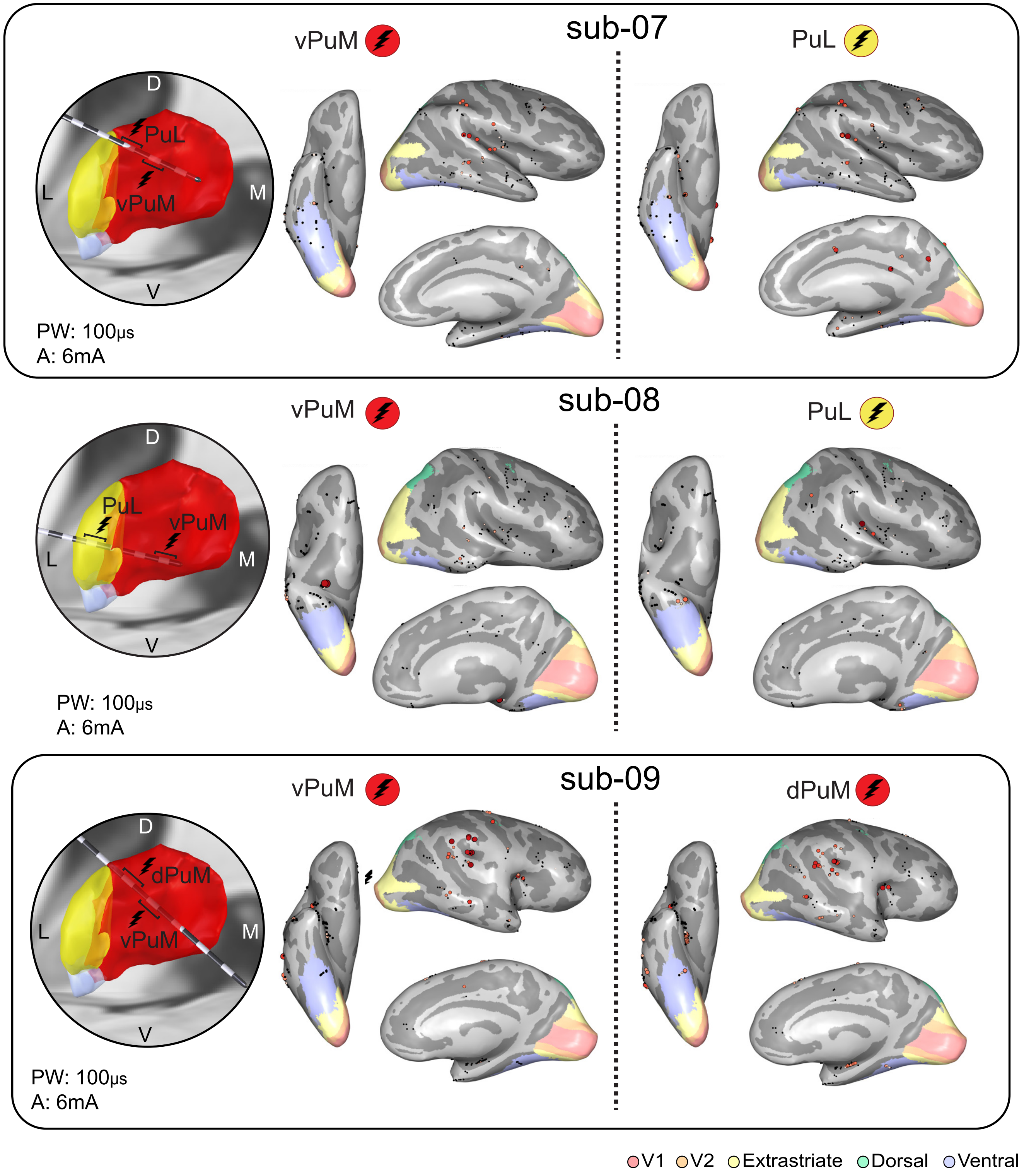

sub-07
vPuM
PuL
D
L
M
V
PuL
vPuM
PW: 100μs
A: 6mA
sub-08
vPuM
PuL
D
L
M
V
PuL
vPuM
PW: 100μs
A: 6mA
sub-09
vPuM
dPuM
D
L
M
V
dPuM
vPuM
PW: 100μs
A: 6mA
V1
V2
Extrastriate
Dorsal
Ventral

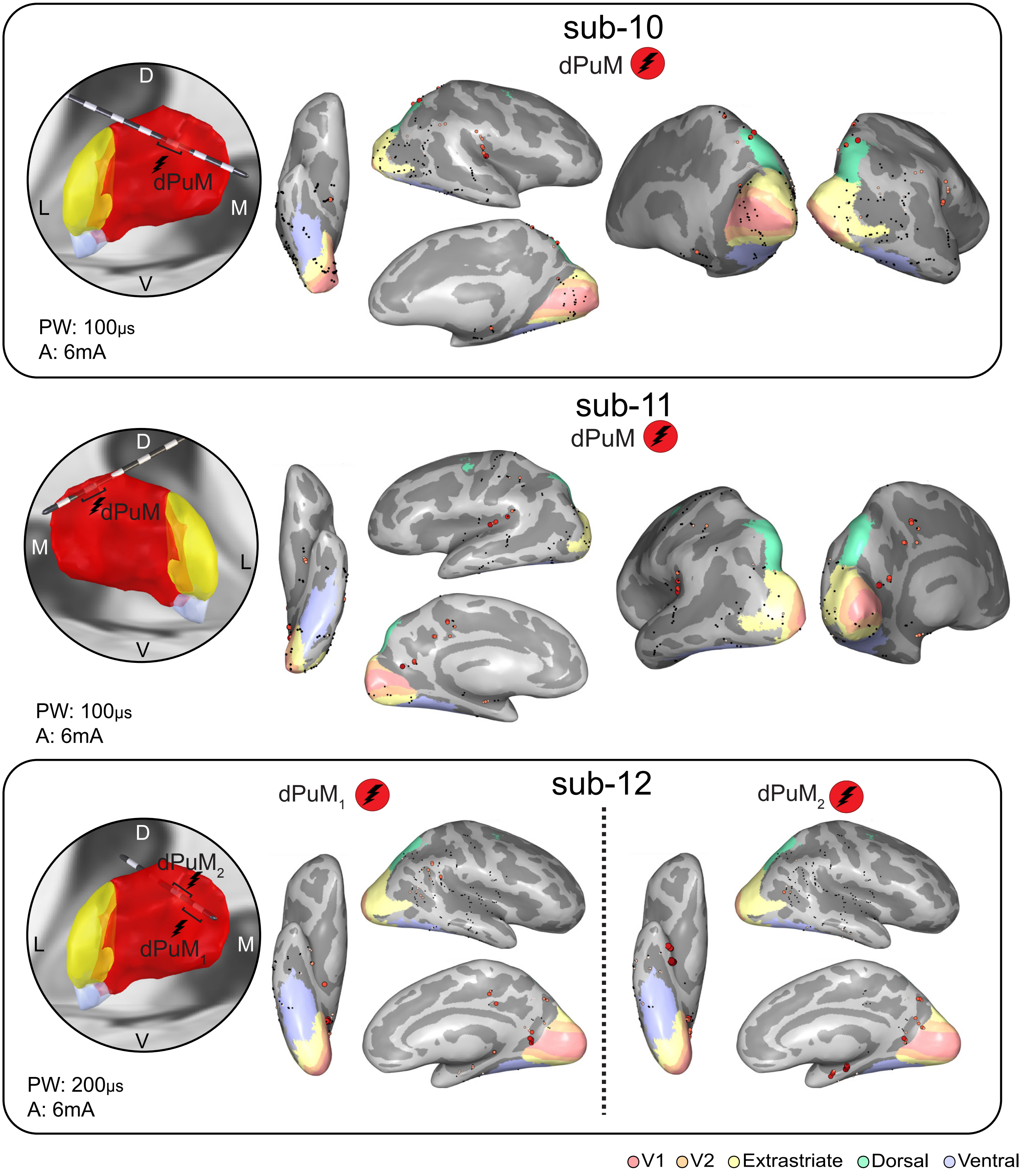
sub-10
dPuM
D
dPuM
L
M
V
PW: 100µs
A: 6mA
sub-11
dPuM
D
dPuM
M
L
V
PW: 100µs
A: 6mA
sub-12
dPuM1
dPuM2
D
dPuM2
L
dPuM1
M
V
PW: 200µs
A: 6mA
V1
V2
Extrastriate
Dorsal
Ventral

**Supplemental Figure 3 Visualization of all subjects pulvinar leads implanted in MNI152 space.** The pulvinar and electrode trajectories were visualized using LeadGroup (LeadDBS 3.0) and the Morel pulvinar atlas. The pulvinar was segmented into three sub-nuclei: medial (red), lateral (yellow), and inferior (grey). Inflated brain surfaces were rendered using a combination of visual atlases: Benson (V1, V2, V3), Rosenke (h0c4v, FG1-4), Wang (TO1-2, LO1-2, V3AB, IPS1-5, SPL1, FEF). The strength of pulvinar-cortical effective connectivity at the most medial and lateral stimulation sites are shown for all subjects.

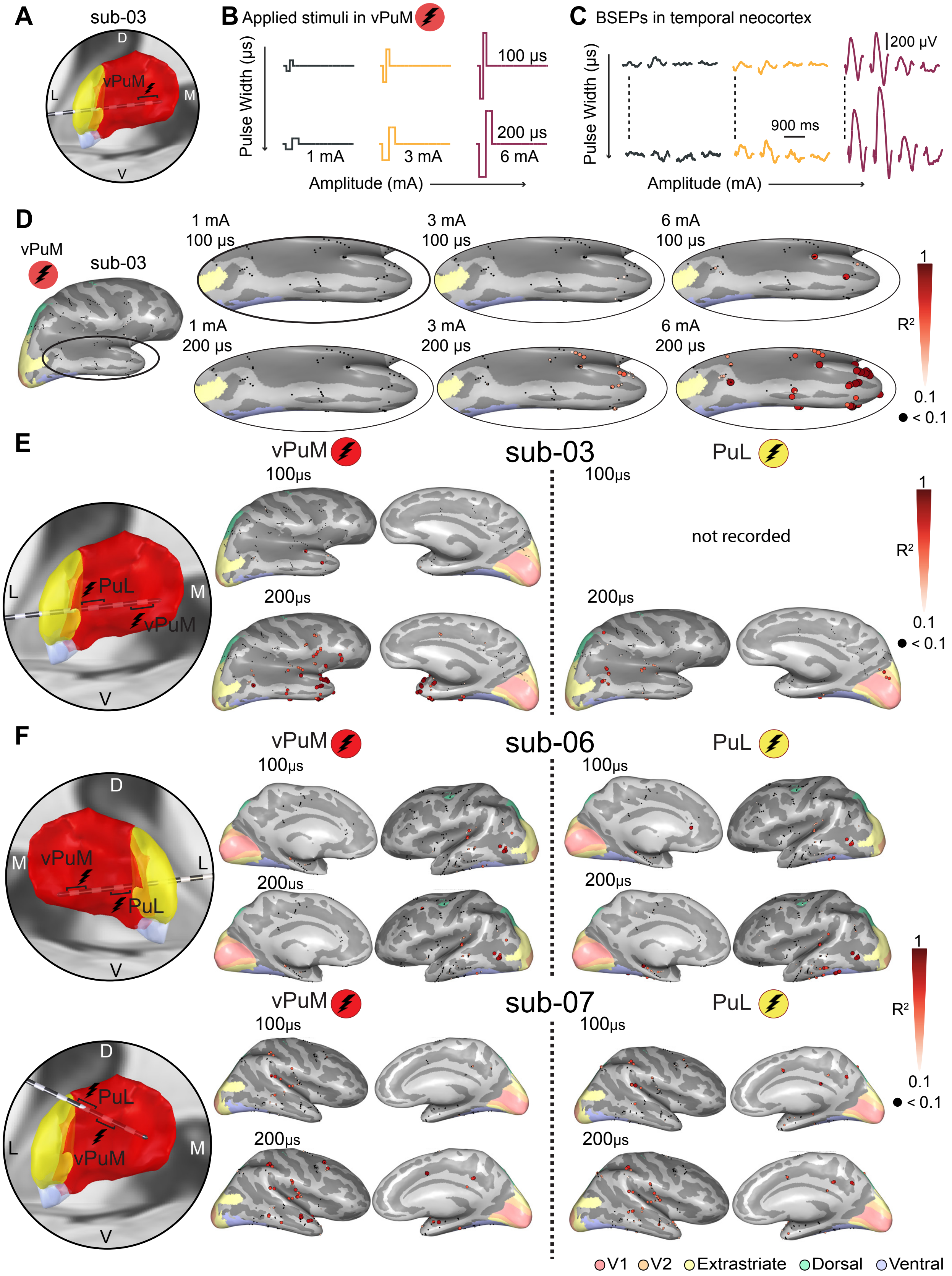
A
sub-03
D
L
M
V
vPuM
B Applied stimuli in vPuM
Pulse Width (µs)
100 µs
200 µs
1 mA
3 mA
6 mA
Amplitude (mA)
C BSEPs in temporal neocortex
200 µV
900 ms
Pulse Width (µs)
Amplitude (mA)
D
vPuM
sub-03
1 mA
100 µs
3 mA
100 µs
6 mA
100 µs
1 mA
200 µs
3 mA
200 µs
6 mA
200 µs
1
R²
0.1
< 0.1
E
vPuM
sub-03
PuL
100µs
100µs
not recorded
200µs
200µs
PuL
vPuM
F
vPuM
sub-06
PuL
100µs
100µs
200µs
200µs
vPuM
sub-07
PuL
100µs
100µs
200µs
200µs
V1
V2
Extrastriate
Dorsal
Ventral

**Supplemental Figure 4 Stimulation condition considerations when stimulating the pulvinar.** **A)** Example subject-03 that underwent pulvinar stimulation of multiple stimulation parameters **B)** Description of the stimulation conditions applied in the medial pulvinar. **C)** Brain stimulation evoked responses (BSEPs) in four measurement electrodes to each stimulation paradigm. The dotted line indicates the first electrode. Amplitude is maximal when delivering a pulse of 6mA and 200 microseconds. **D)** Strength of pulvinar-cortical effective connectivity in the lateral temporal cortex when stimulating the medial pulvinar at differing stimulation parameters. **E)** Pulvinar-cortical effective connectivity mapping is consistent when stimulating both at 100 and 200 microseconds in subject-03 **F)** Pulvinar-cortical effective connectivity mapping causes more variable effects when stimulating both at 100 and 200 microseconds in subject-06 and subject-07. Both E and F have pulses of amplitude 6mA. Both subject-06 and subject-07 had leads near the border of the pulvinar, which may explain inconsistent outputs when increasing the current delivered.

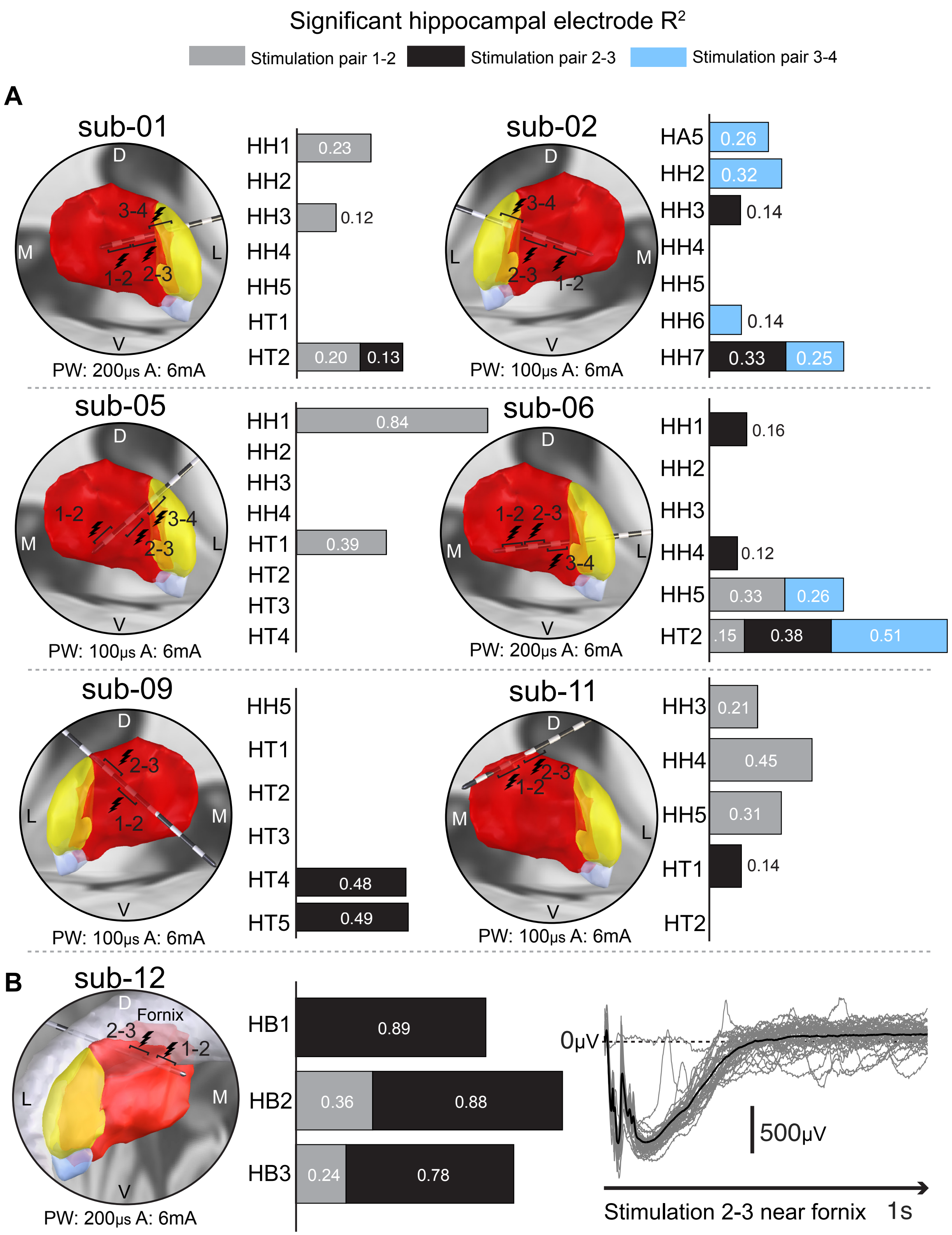

Significant hippocampal electrode $R^2$
Stimulation pair 1-2
Stimulation pair 2-3
Stimulation pair 3-4
A
sub-01
D
M
L
V
3-4
1-2
2-3
PW: 200µs A: 6mA
HH1 0.23
HH2
HH3 0.12
HH4
HH5
HT1
HT2 0.20 0.13
sub-02
PW: 100µs A: 6mA
HA5 0.26
HH2 0.32
HH3 0.14
HH4
HH5
HH6 0.14
HH7 0.33 0.25
sub-05
PW: 100µs A: 6mA
HH1 0.84
HH2
HH3
HH4
HT1 0.39
HT2
HT3
HT4
sub-06
PW: 200µs A: 6mA
HH1 0.16
HH2
HH3
HH4 0.12
HH5 0.33 0.26
HT2 .15 0.38 0.51
sub-09
PW: 100µs A: 6mA
HH5
HT1
HT2
HT3
HT4 0.48
HT5 0.49
sub-11
PW: 100µs A: 6mA
HH3 0.21
HH4 0.45
HH5 0.31
HT1 0.14
HT2
B
sub-12
Fornix
PW: 200µs A: 6mA
HB1 0.89
HB2 0.36 0.88
HB3 0.24 0.78
0µV
500µV
Stimulation 2-3 near fornix
1s

**Supplemental Figure 5 Stimulation responses from pulvinar to hippocampus. A)** Each subject with significant pulvinar-hippocampal effective connectivity is shown below (7 total). We plot a stacked bar chart of their coefficient of determination ($R^2$) for each stimulation-electrode contact pair. Only responsive stimulation-electrode contact pairs are shown. The coefficient of determination is relatively weak and variable with no clear effective connectivity map. Sub-06 showed no hippocampal stimulation response at 100 microseconds but did at 200 microseconds **B)** Stacked bar chart as before in A. However, stimulation contacts approach the fornix, causing highly significant brain stimulation evoked responses (gray=all trials, black line=mean response). MRI shows electrode positions approaching the fornix. Electrode targets are as follows: HH=hippocampus head, HB=body, HT=tail, HA=hippocampus/amygdala where, # represents the contact number on the lead. The Fornix was visualized using the Fornix FMRIB FA template in LeadDBS.[11]
